# Virion-associated influenza hemagglutinin clusters upon sialic acid binding visualized by cryo-electron tomography

**DOI:** 10.1101/2024.10.15.618557

**Authors:** Qiuyu J. Huang, Ryan Kim, Kangkang Song, Nikolaus Grigorieff, James B. Munro, Celia A. Schiffer, Mohan Somasundaran

## Abstract

Influenza viruses are enveloped, negative sense single-stranded RNA viruses covered in a dense layer of glycoproteins. Hemagglutinin (HA) accounts for 80-90% of influenza glycoprotein and plays a role in host cell binding and membrane fusion. While previous studies have characterized structures of receptor-free and receptor-bound HA in vitro, the effect of receptor binding on HA organization and structure on virions remains unknown. Here, we used cryo-electron tomography (cryoET) to visualize influenza virions bound to a sialic acid receptor mimic. Overall, receptor binding did not result in significant changes in viral morphology; however, we observed rearrangements of HA trimer organization and orientation. Compared to the even inter-glycoprotein spacing of unliganded HA trimers, receptor binding promotes HA trimer clustering and formation of a triplet of trimers. Subtomogram averaging and refinement yielded 8-10 Å reconstructions that allowed us to visualize specific contacts between HAs from neighboring trimers and identify molecular features that mediate clustering. Taken together, we present new structural evidence that receptor binding triggers clustering of HA trimers, revealing an additional layer of HA dynamics and plasticity.

## INTRODUCTION

Influenza viruses are highly contagious respiratory pathogens that cause approximately 1 billion cases of flu in humans each year. Influenza initiates infection through receptor-mediated endocytosis, where the trimeric hemagglutinin (HA) glycoprotein binds sialic acid containing host cellular receptors (1). Individual interactions between HA and sialic acid are weak; polyvalent interactions between multiple simultaneous HA-sialic acid binding events are required for successful viral entry (2).

HA is a 200-kDa protein that is structurally divided into the globular domain (HA1), which includes the sialic acid receptor binding site, and the HA2 domain, which contains the fusion peptide, a hydrophobic stretch of amino acids that inserts into the endosomal membrane during fusion with the viral envelope (3). Structural studies of the soluble HA ectodomain revealed that sialic acid binding does not induce large structural rearrangements in HA ectodomains (4-6). In contrast, biophysical analyses of full-length HA on the virion surface indicated that sialic acid binding induces movement of the fusion peptide (7). This observation emphasizes that ectodomain structures provide only a limited amount of insight into the dynamic processes of HA-receptor interaction, and a full understanding of the HA-mediated virus binding and entry into cell requires structural analysis of HA on intact virions. Previous cryo-electron tomography (cryoET) studies, including our work, have shown that spherical influenza virions carry 300 to 500 glycoproteins, with consistent inter-glycoprotein spacing of between 9-11 nm (8-10). Moreover, biophysical studies showed that hemifusion with the target membrane requires cooperativity among three to four neighbouring HA trimers (11, 12), suggesting that HA trimers form inter-trimer interactions to facilitate the process. In this context, receptor binding may promote HA cooperativity and clustering. However, the effects of receptor binding on HA structure in the context of virions have not previously been investigated.

In the current study, we employed cryoET for ultrastructural studies of the A/Puerto Rico/8/34 (H1N1) influenza A virus (henceforth referred to as PR8) incubated with a sialic acid-containing receptor mimic. While receptor binding did not result in significant changes in viral morphology, we observed formation of HA clusters composed of triplet of trimers in a receptor-binding dependent manner. Within this assembly, HAs form direct interactions with HA protomers of neighboring trimers, using both polar and electrostatic interactions. Our study reveals that receptor-bound HA trimers are highly mobile, suggesting that disrupting either HA mobility or clustering may offer a new strategy for targeting influenza virus.

## RESULTS

### CryoET of receptor-bound influenza virus reveals little changes in viral morphology

To evaluate whether receptor binding causes structural rearrangements in HA or changes in viral morphology, we obtained tilt-series of PR8 with and without the receptor mimic sialylneolacto-N-tetraose c (LSTc). LSTc is a glycan containing a terminal α2–6-linked sialic acid; it productively binds and inhibits PR8 and other influenza A virus strains (13). Prior to plunge freezing, influenza virions were incubated with 100 μM or 6.5 mM of LSTc for at least 15 minutes to reach equilibration (Table S1).

Tomogram reconstruction of PR8 and PR8-LSTc allowed for clear identification of virus components (Fig 1, Movie S1-2). Similar to previous cryoET studies at neutral pH (8, 14, 15), PR8 virions are membranous viruses of spherical or oval morphology. Beneath the lipid bilayer, most virions showed clear density for the matrix protein (M1) layer, as well as the viral ribonucleoprotein (vRNP) complex. We were also able to discern the two distinct glycoproteins, HA and NA, based on the clear difference in their size and shape with HA appearing taller and cylindrical (Fig 1A, B). To quantify potential morphological differences between receptor-free and receptor-bound virions, we applied the convolutional neural network (CNN) analysis pipeline we previously developed (Fig. S1) (8). Consistent with our recent findings, influenza virions were extremely pleomorphic (Fig 1C, D). The median long and short axes measurements were 124 nm and 106 nm for receptor-free, and 129 nm and 108 nm for receptor-bound virions, with the median axial ratio of both populations of 1.1 (Table S3). Therefore, we concluded that receptor binding does not result in large morphological changes in influenza virions.

**Figure 1.**
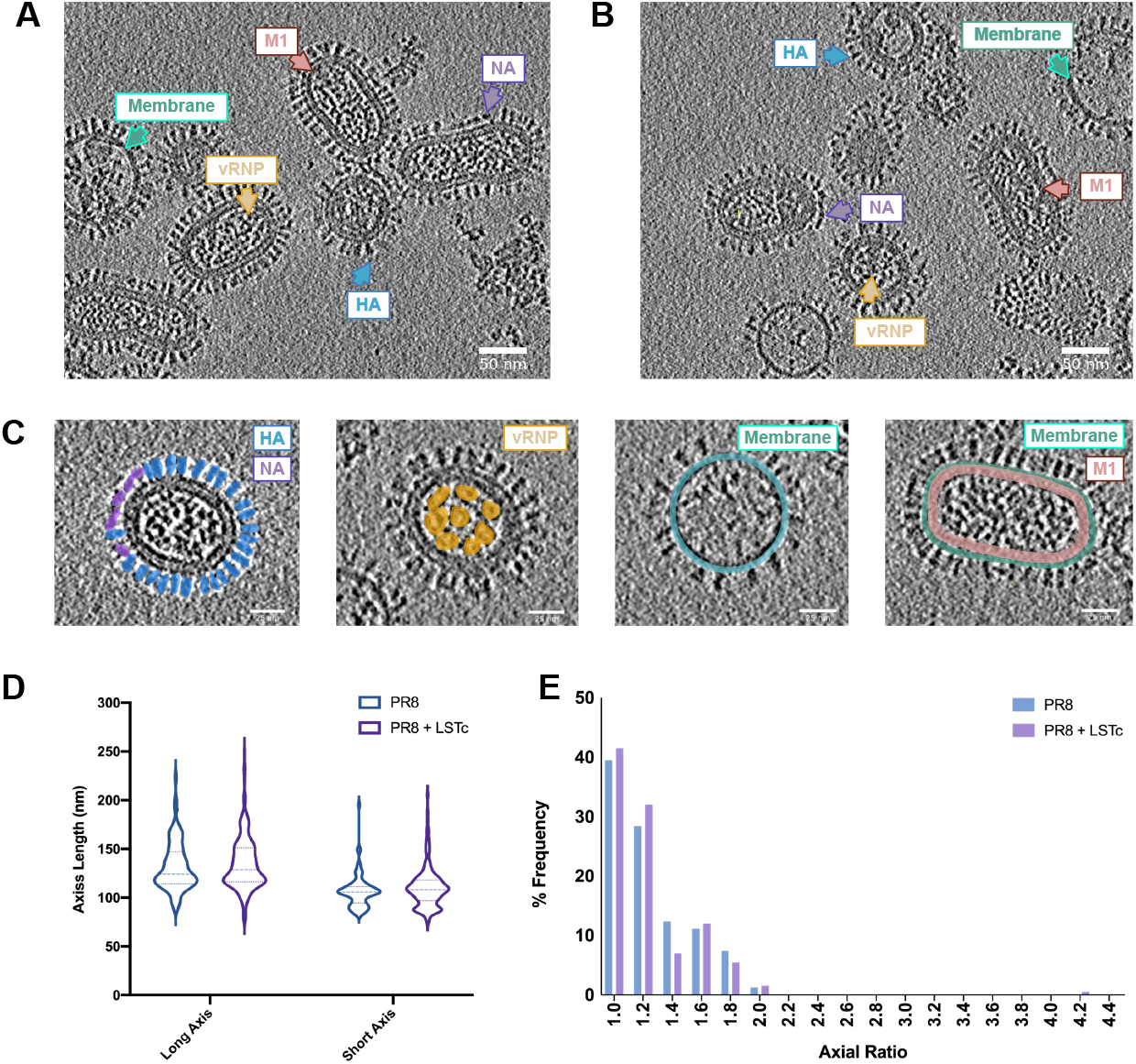
Cryo-electron tomography reveals little morphological difference between influenza virions incubated with soluble receptor mimic and influenza virions alone. **A, B**. Central slice through representative tomogram of PR8 virions (**A**) or of PR8 virions with 100 μm LSTc (**B**). Scale bars = 50 nm. **C**. Violin plot for long and short axis measurements of PR8 virions with and without LSTc. Middle dotted line indicates the median value; top and bottom dotted lines indicate interquartile range. Width of the plot correlates with the number of particles at that value.**D**. Histogram of axial ratios of PR8 virions.

### Receptor binding promotes asymmetrical clustering of neighbouring HA trimers

To investigate dynamic changes in HA ordering and structure upon receptor binding, we conducted reference-free subtomogram averaging on receptor-free and LSTc-bound HAs. We analyzed virions where both the M1 layer and the vRNP density were present, as influenza virions missing either component would be less functionally relevant. Since influenza glycoproteins are perpendicular to the viral membrane, initial Euler angles can be estimated using the surface normal of HA coordinates picked using the CNN pipeline described in Fig S1 (Fig S2). These coordinates and angles were used to extract bin4 subtomograms using Warp and subjected to subtomogram refinement within RELION 4 with C1 symmetry (16, 17). Several rounds of 2D classification were carried out to discard outliers and junk particles (Fig S3-4). Both HA and HA+LSTc reconstructions revealed a glycoprotein array, where clear density was shown for a patch of 7-10 HA trimers and the viral membrane (Fig 2). At this resolution, the distinct three-fold symmetry had only emerged for the central HA of the unliganded HA reconstruction (Fig 2A, B). Peripheral HAs appeared as cylindrical density and their three protomers were indistinguishable (Fig 2A, B). This observation contrasted with LSTc-bound HA, where distinct three-fold symmetry was shown for not only the central HA but two more HAs on either side (Fig 2C, D). Thus, receptor binding induced rearrangement on the viral surface with neighbouring HAs oriented towards each other to form a triplet of HA trimers (Fig 2C, D).

**Figure 2.**
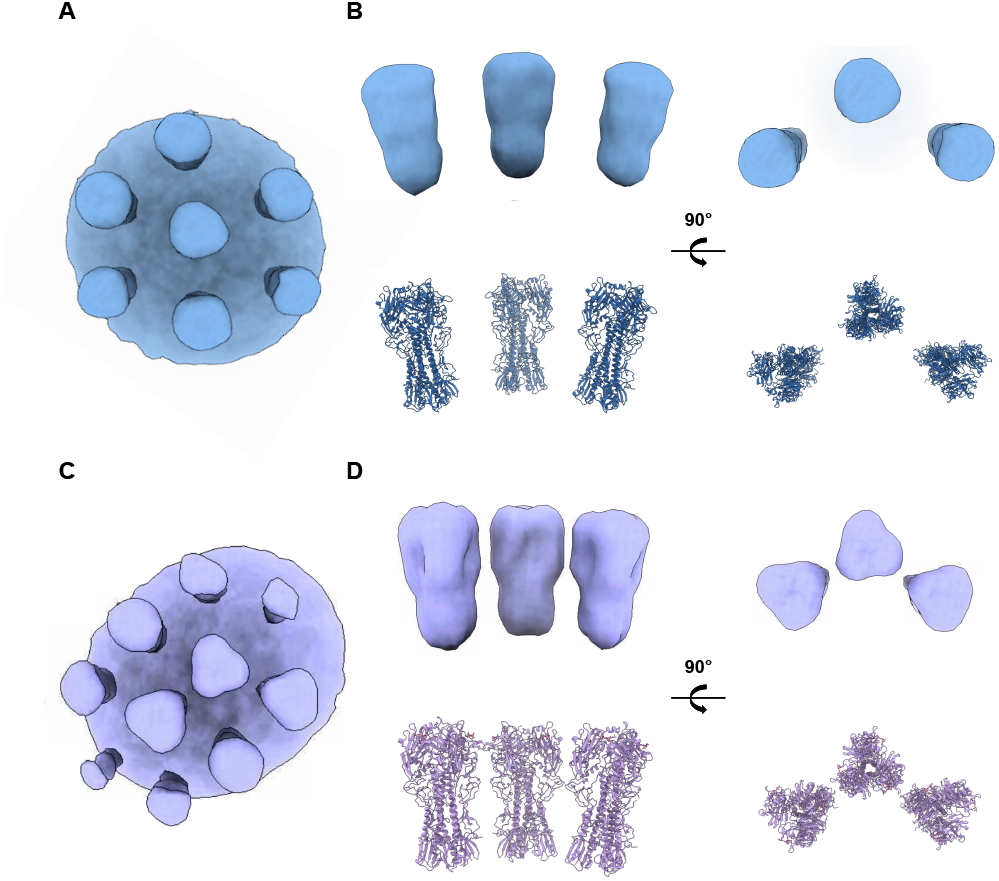
Subtomogram averaging of HA reveals altered glycoprotein organization upon receptor recognition. **(A)** Top view of a bin4 reconstruction depicting a HA patch from PR8; reconstruction is looking down at a patch of HA glycoproteins atop the viral membrane. Untreated influenza has regular organization of HAs whereas LSTc-bound influenza has clustered HAs. **(B)** Zoom in view of three central HAs from PR8 virions. (Upper) The side and top views are presented. (Lower) Crystal structures of PR8 HA (PDB: 1RU7) fitted into the reconstruction. **(C)** Top view of a bin4 reconstruction depicting a HA patch from PR8 virions incubated with LSTc; reconstruction is looking down at a patch of HA glycoproteins atop the viral membrane. **(D)** Zoom in view of three central HAs from PR8 virions incubated with LSTc. (Upper) The side and top views are presented. (Lower) Crystal structures of PR8 HA bound to LSTc (PDB: 1RVZ) fitted into the reconstruction. Terminal sialic acid is shown as sticks. PR8 reconstructions and models are shown in light blue and PR8 + LSTc reconstructions and models are shown in lavender.

To characterize this arrangement of HA trimers further, we calculated the inter-HA distance between the central HA and its two closest neighbours in both receptor-free and receptor-bound structures (Fig S5). Receptor-free HA had a narrow distribution of inter-HA spacing, with the median distance between the central HA and its two closest neighbours between 84 Å and 91 Å, with an interquartile range of 80-87 Å and 87-95 Å respectively. The distribution of inter-HA distances in receptor-bound HA was wider, with interquartile range of 75-90 Å and 77-99 Å. We also saw a general decrease in the spacing between the three central HAs, with the central HA on average 4 Å and 7 Å closer to its two neighbours. Upon 2D classification of the HA trimer array, we observed classes in which the central HA pivoted closer to either adjacent HA when compared with the consensus reconstruction (Fig S6). This variance is likely caused by dynamic HA rearrangements upon receptor binding, suggesting that instead of inducing structural rearrangements within individual protomers or within the HA trimer, receptor binding induces HA clustering.

### Subnanometer resolution subtomogram average of PR8 HA reveals domain organization of membrane-bound HA

From the low-resolution HA array, we continued refinement for the central HA trimer with C1 symmetry. Aligned subtomograms were gradually unbinned and local refinement with 2D classification steps were conducted using a focused mask in RELION (Fig S3) (17). Lastly, pose refinement was conducted in M (18), yielding an HA map at a global resolution of 8.9 Å from a set of 12,063 particles (Fig 3A, Fig S7). At this resolution, both isosurface and projection slices clearly show separation between individual HA protomers and individual helices within HA2 (Fig 3A, C). Furthermore, we could observe density corresponding to the viral membrane at lower contours (Fig 3A). Our reconstruction resembles reported HA structures (4, 5, 8, 19, 20), showing a trimeric protein with three-fold symmetry and of approximately 14 nm in height and 8 nm in width. However, flexible fitting of a PR8 HA ectodomain crystal structure into the map showed subtle relaxation of the head domain and constriction of the stem, with a global RMSD of 1.89 Å (Movie S3). Likely, the constriction of the stem domain corresponds to tethering from the transmembrane helices and cytoplasmic tail that is not present in the crystal structure. Taken together, these data represent the highest resolution view of virion-bound HA obtained by cryoET to date.

**Figure 3.**
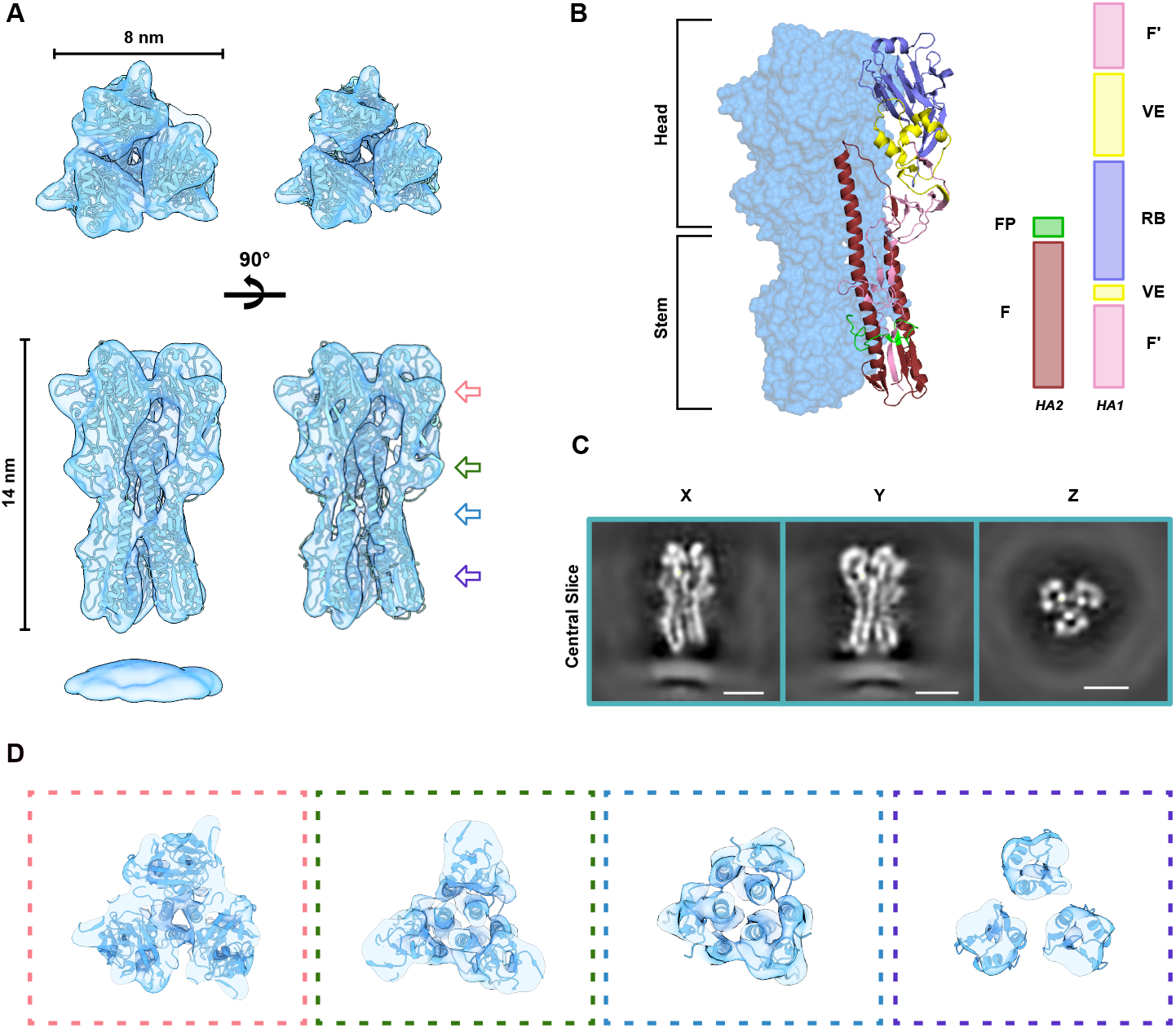
Subtomogram average of PR8 HA. **(A)** Top and side view of a cryoET map of the HA trimer at two contour levels flexibly fitted with a PR8 HA ectodomain crystal structure (PDB: 1RU7) (*29, 30*). (**B)** Architecture of flexibly fitted HA. Two protomers are shown as blue surfaces and the third protomer as cartoon. Individual HA domains are colored as following: fusion [HA1] (F’) – pink; vestigial esterase (VE) – yellow; receptor binding domain (RB) – purple; fusion peptide (FP) – green, and fusion (F). (**C)** Central projection slice through the masked HA map viewed in X, Y, and Z directions. Scale bars are 5 nm. **(D)** Clipped views through the HA reconstruction. Red box depicts RB, green box depicts VE, and two views from the HA stem are shown in blue and purple boxes.

### Neighbouring HA trimers form a HA1-HA1 interface orchestrated by mirrored ionic interactions

We subsequently investigated the configuration of trimers seen in LSTc-bound samples using subtomogram averaging of influenza virions incubated with 100 μM LSTc and 6.5 mM LSTc (Fig 4). Refined coordinates from the bin4 reconstruction (Fig 2C, D) were used to extract bin2 particles. For both LSTc-bound samples, the triplet of HA trimers we described above persisted at higher resolution reconstructions (Fig 4A). We performed three independent refinements for the sample with 6.5 mM LSTc using central HA alone, central HA and the closest HA partner (pair I), and central HA plus the second closest HA partner (pair II).

**Figure 5.**
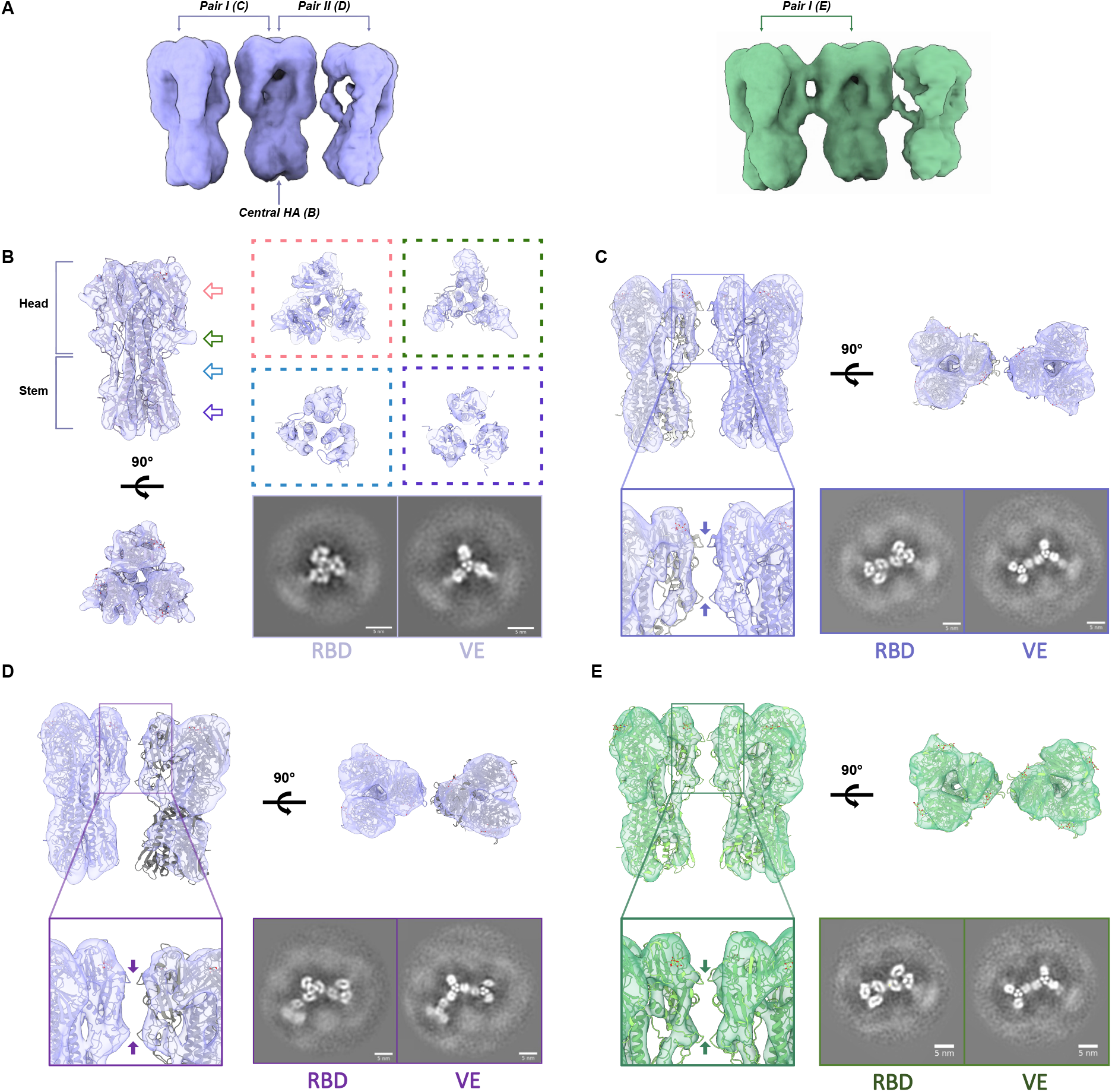
Receptor binding induces inter-HA contacts but not intra-HA domain rearrangement. **(A)** Reconstruction of HA trimer of trimers incubated with (left) 6.5 mM LSTc or (right) 100 μM LSTc. Maps were obtained using bin2 particles. **(B)** (Left) Top and side views of the central HA in the trimer of HAs Pair I in the trimer of HAs. (Upper right) Clipped views through the HA reconstruction. Red box depicts RB, green box depicts VE, and two views from the HA stem are shown in blue and purple boxes. (Lower right) Central projection slice through the HA map at the RB and VE domains. (**C-E**). (**C**) Pair I in the trimer of HAs. (**D**). Pair II in the trimer of HAs. (**E**) Pair I in the trimer of HAs from virions incubated with 100 μM LSTc. (Upper) Side and top views of each map were flexibly fitted with two copies of a PR8 HA ectodomain crystal structure bound to LSTc (PDB: 1RVZ) (*29, 30*). (Inset) zoomed in view of RB and VE interfaces highlighted by arrows. (Lower right) Central projection slice through each reconstruction at the RB and VE domains. All scale bars are 5 nm.

Subtomogram averaging of the central HA reached a global resolution of 8.5 Å with 17,266 particles (Fig S7). Both HA1 and HA2 domains were well defined at this resolution, as well as secondary structure elements, especially in HA1 (Fig 4B). Similar to the receptor-free HA reconstruction, we were able to trace α-helices at this subnanometer resolution. While we saw similar relaxation in the head domain and constriction within the stem domain as in the receptor-free HA, there were no additional domain rearrangements we could detect within receptor-bound HA.

To further refine our particle set, we conducted another round of 2D classification to remove approximately 25% of particles where the two neighbouring HAs were further away from the central HA (Fig S4). The remaining 12,379 particles were centered on pair I and subjected to refinement, with the final map at C1 symmetry reaching 8.9 Å resolution (Fig S7). Inspecting the reconstruction for pair I, we confirmed the appearance of a HA1-HA1 interface between the two HA trimers (Fig 4C). The two HA molecules were mirror-image of one another, with the head domain of one protomer coming in proximity with the adjacent HA head. Between the two HAs, the flexible loop of the two receptor-binding regions and the vestigial esterase loops were anti-parallel to each other (Fig 4C). Moreover, our reconstruction of pair I from virus incubated with 100 μM LSTc was nearly identical (Fig 4E). For both maps, density was weaker in the stem region of the protomer facing the central HA, indicating possible flexibility in the region.

We observed a similar interface for pair II (Fig 4D). The final reconstruction contained a set of 10,548 particles and reached a resolution of 10 Å (Fig S4). Density for the adjacent HA was weaker in comparison to the adjacent HA of pair I, and this weaker density was represented by its lower local resolution (Fig S4). Likely, there was more flexibility in the adjacent HA as the inter-HA distances also showed the adjacent HA in pair 1 was closer than that of pair 2 (Fig S5). However, it was apparent that the two neighbour-facing protomers of the central HA engaged its opposing domains similarly, with the globular heads packing against each other (Fig 4C, E). Taken together, we discovered the formation of a new inter-HA interface arrangement that could enhance efficient receptor-binding and cell entry.

### Molecular details of interactions that mediate clustering of receptor-bound HA

Lastly, we investigated the interaction interfaces of receptor-bound HA (Fig 5). To define HA-HA interfaces, we flexibly refined crystal structures of LSTc-bound HA within our reconstructions, and the refined models underwent energy minimization to determine optimal contacts. The two interfaces were both located within HA1, one in the receptor-binding domain and the second at the end of the vestigial esterase domain. Residing just outside of the receptor-binding site and implicated in neutralizing antibody interactions (21), the 140-loop constitute the first interface (Fig 5B). Glu142 is the main driver of this network, forming a salt bridge with Arg149’ and hydrogen bonds with His141’ and its mirroring residue, Glu142’ (Fig 5C). Within the lower interface, His276 forms hydrogen bonds with Glu277’ of the opposing protomer, and vice versa (Fig 5D). Altogether, our subtomogram averages provide strong evidence that these polar and electrostatic interactions drive interface contacts between adjacent HA trimers, and help promote the stable triplet of trimer HA configuration upon receptor binding.

**Figure 5.**
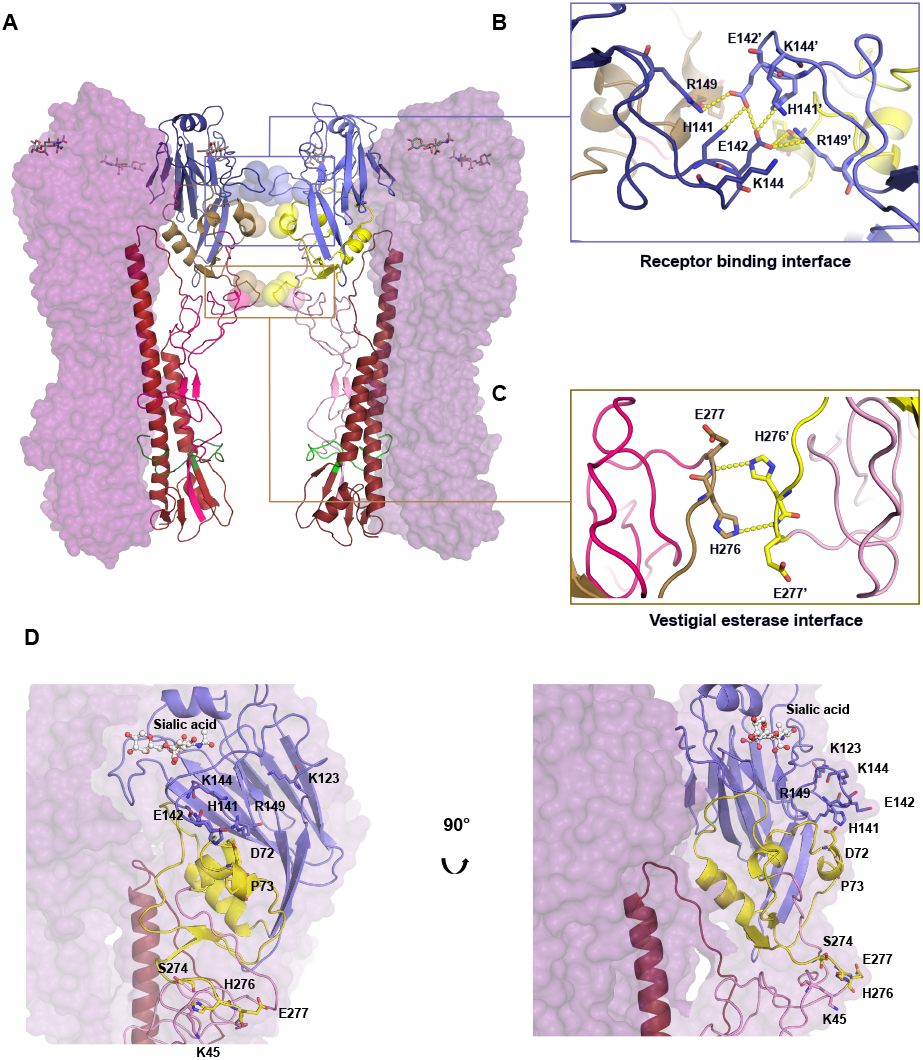
HA-HA interface reveals mirror image contacts within HA1. (**A**) Model of interacting HA pair. Two protomers from each HA trimer are represented as a purple surface and the third as colour-coded cartoon. Terminal sialic acids are represented as sticks. Domains are coloured as according to the key in **Figure 3**. Residues within 5 Å of the neighbouring protomer are shown as spheres. (**B**) Zoomed in view of receptor binding domain interactions. Hydrogen and ionic bonds are shown as yellow dashed lines. (**C**) Zoomed in view of vestigial esterase domain interactions. Hydrogen and ionic bonds are shown as yellow dashed lines. (**D**) Zoomed in view of proximal residues indicated as spheres in (**A**). Terminal sialic acids are shown as stick and balls.

**Figure 6.**
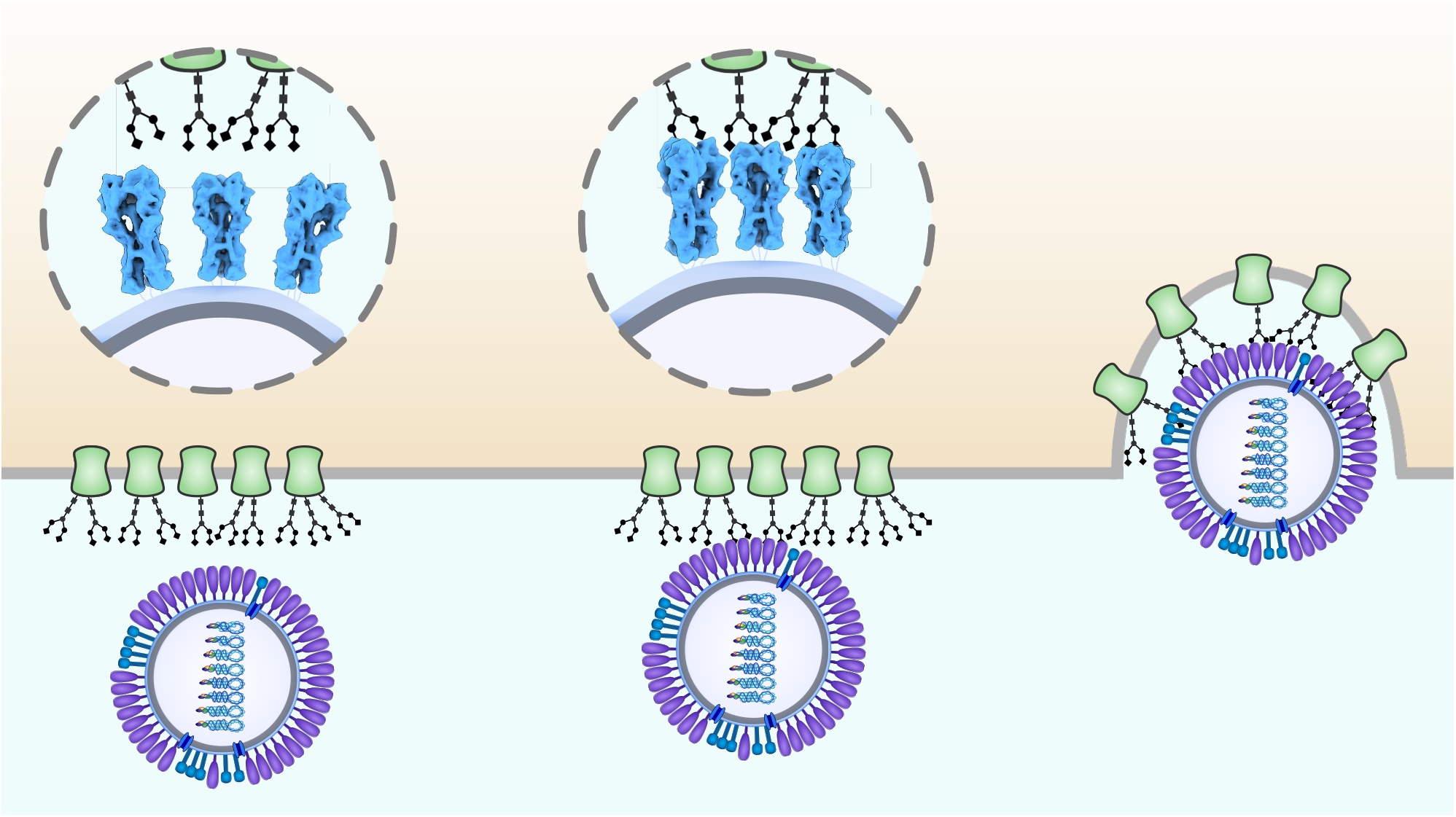
Model of influenza HA binding and entry. Influenza virions travel within the respiratory tract prior to cellular entry. During this stage, HAs are capable of movement as the virions tumble to better sense the environment. Upon contact with the sialylated cell surface, local coordination of neighbouring HAs allow for multivalent interaction with the cell within the contact region to induce productive cellular entry.

## DISCUSSION

In summary, our study uses cryoET to illustrate receptor binding to native virions, which is an initial step of influenza virus infection. Through tomographic analysis and subtomogram averaging, we observed rearrangement of the HA glycoprotein array, resulting in HA clustering to form a triplet of trimers. We also characterized a previously undetected HA-HA interface that emerges upon receptor binding. These analyses gave rise to several subnanometer resolution subtomogram averages of influenza HA, including: 1) the highest reported resolution of a virion-bound HA subtomographic reconstruction and 2) the first elucidation of the structure of virion-bound HA in the presence of a cellular receptor mimic.

Despite its pleomorphism, influenza virions exposed to a sialic acid containing receptor mimic did not exhibit any morphological changes in the virion, consistent with previous studies (22-24). However, we observed changes in HA organization on the viral surface upon receptor binding, whereby receptor binding brought HA trimers closer together to form a triplet of trimers. We also discovered upon receptor binding a new interface formed between HA protomers in neighboring HA trimers as they cluster. This receptor-triggered clustering is likely functionally relevant. Previous studies have established that influenza virions bind cell surface sialic acid containing cellular receptors multivalently (2, 25-27). This multivalent attachment increases the possibility that clustering of neighbouring HAs would further enhance binding by allowing the virion to bind to an entire patch of surface glycans. Thus, we propose that once receptor-bound, dynamic HA movements become stabilized and form steady interactions with adjacent trimers to prepare for membrane fusion. We do not, however, observe large domain-level rearrangements within singular HA trimers, which is consistent with previous crystallographic studies (5). However, this process may be accompanied by smaller secondary structure and local changes, as these types of changes are beyond the resolution of the current study. Nevertheless, we visualized highly packed HA trimers are mobile on the virion surface and impact the structure and organization of neighbouring glycoproteins,, as was previously proposed but not observed (28). It is likely that, given HA’s millimolar affinity for sialic acid, HA clustering in presence of its receptor enhances multivalent binding to productively achieve cellular entry. Thus, even local changes, such as receptor binding initially on one HA, could cause a rippling avidity effect through the viral surface to promote entry into the cell.

Interestingly, coordination between neighbouring HAs is relevant in the context of other processes beyond receptor binding. Over the past two decades, approximately three HA molecules have been shown by kinetics to form a local fusion cluster that are required for fusion between endosomal and viral membranes (11, 12). During acidification in an infected endosome, one HA can start the fusion process via insertion into the target membrane, but membrane fusion occurs rapidly only after 2-3 additional proximal HA molecules enter hemifusion. Thus, our full viral reconstructions corroborate and support the formation of a fusion cluster during influenza infection by a receptor-bound HA cluster consisting of three adjacent HA trimers on the virion surface tilting towards each other and thereby provide additional insights into the influenza entry pathway.

Our study further demonstrates the intricacies of the influenza entry pathway and shows additional dynamics of HA during receptor binding. We characterized the reorganization of glycoprotein packing on the virion surface upon sialic acid engagement, and revealed a previously unseen formation of a triplet of HAs. Through subtomogram averaging of influenza HA, we reconstructed the highest resolution virion-bound HA reported, and the first receptor-bound HA reconstructions from whole virions. These maps allowed us to describe a novel interface between adjacent HAs that is established post receptor binding. In conclusion, this report provides the first structural view of the intact, receptor-bound influenza virions, and provides further evidence that HAs orchestrate a coordinated mechanism for optimal viral entry. We anticipate that better understanding of HA cooperativity will advance our understanding of the molecular determinants of a successful infection and illuminate additional strategies for therapeutic development.

## MATERIALS AND METHODS

### Virus and reagents

A/PR/8/34 (PR8) influenza virus was grown in embryonated chicken eggs and purchased from Charles River Laboratories (Now AVS Bio. Catalog #: 10100374) and stored at -80 °C before use. LS-Tetrasaccharide c (LSTc) was purchased from Dextra (Catalogue #: SLN506) and stored at -20 °C before use.

### Cryo-ET sample preparation and tilt-series data collection

PR8 influenza virus was diluted to 1 mg/mL or 0.5 mg/mL, and receptor-bound samples were incubated with 100 µM or 6.5 mM final LSTc concentration. Virus was mixed with BSA-coated 10 nm colloidal gold (Sigma-Aldrich) and 3 µL were applied to glow-discharged R2/2 holey carbon copper grids (Quantifoil). These grids were manually back blotted for 2 to 3s with filter paper, and rapidly frozen in a liquid ethane/propane mixture using an EMS-002 rapid immersion freezer. Prior to imaging, vitrified grids were stored in LN2. Grids were imaged on a Thermo Fisher Scientific Titan Krios electron microscope operating at 300 kV with a K3 camera (Gatan, Pleasanton, USA) and a Gatan energy filter at a slit width of 20 eV centered on the zero-loss position. Tilt series of PR8 alone and PR8 with 100 µM LSTc were recorded from 0 to ±66° in 2-degree increments at a magnification of 42,000x (corresponding to a calibrated 2.0873 Å of pixel size) using a dose symmetric scheme (31). Raw movies of these samples were acquired at an electron dose of 1.86e-/Å2. Tilt series of PR8 incubated with 6.5 mM LSTc were recorded from 0 to ±51° in 3-degree increments at the same magnification. Raw movies of this sample were acquired at an electron dose of 3e-/Å2. The sample was imaged in counting mode at each tilt angle as movies (4-6 frames), and the accumulated dose of all movies was 120 e-/Å2. The defocus range of all tilt series is 4-8 µm. Data were acquired automatically under a low-dose mode using the SerialEM software (32).

### Tilt-series preprocessing and tomogram generation

Raw movies were imported into Warp (v1.1beta) (16), where movie frame alignment and CTF estimation were carried out. Dose-weighted tilt series were exported from Warp and imported into IMOD for batched fiducial-based tilt series alignment (33). Aligned stacks were reimported into Warp for tomogram reconstruction at bin4 (8.35 Å/pixel).

### Tomographic analysis

Tomograms were imported into EMAN2.99 and preprocessed to enhance features for subsequent convolutional neural network-based segmentation and particle picking (34). Post-training, HA particle coordinates from each tomogram were saved and modelled as 3D point clouds using the Open3D library (35). Outlier points were removed from point clouds based on the standard deviation of a point’s distance from its closest neighbours from the average distances across the point clouds; This step ideally removes misannotated particles. The agglomerative clustering algorithm implemented in the Scikit-Learn library (36) were used to generate single virion point clouds which were then exported as STL surfaces for automated virion morphology calculations in UCSF Chimera (37).

To filter out virions that did not contain the M1 protein, HA and M1 coordinates were imported as point clouds. Proximity to M1 was used as a metric to conduct distance-based filtering of HA points, and HA coordinates more than 10 nm away from its nearest M1 coordinate were discarded for subtomogram averaging. Lastly, single virion point clouds were imported into a Dynamo v1.15 environment within MATLAB R2023a for initial Euler angle estimation (38). The surface normal for each HA coordinate was calculated and its equivalent rotation matrix was converted into Euler angles. To generate Warp compatible metadata, Euler angles were converted from Dynamo to RELION convention using the eulerangle library and were written into a star file with its corresponding coordinate using the starfile library (39).

### Subtomogram averaging

Initially, HA subtomograms were extracted at a pixel spacing of 8.35 Å and a box size of 48 pixels. This box is large enough to encompass a 3×3 array of membrane-embedded influenza glycoproteins. An initial subtomogram average was generated through reference-free subtomogram averaging of a subset of 1000 subtomograms in RELION 4.0 (17). This average was used as reference for subtomogram alignment of the entire dataset. After several rounds of 3D autorefine in RELION 4.0, 2D classification was carried out on aligned subtomograms to further identify outliers and non-aligned particles. This 2D classification step uses project images obtained from subtomograms, which represent a much faster way to conduct classification in comparison to 3D classification. These particles and duplicate particles were subsequently discarded. This subtomogram averaging scheme was carried out for subtomograms extracted at 4.175 Å/pixel and 2.8 Å/pixel, with more restricted angular sampling. After subtomogram averaging converged at 2.8 Å/pixel, subtomogram coordinates were imported into M v1.1beta and three rounds of particle pose refinement were carried out (18).

Lastly, subtomograms were exported at 2.1 or 2.2 Å/pixel and aligned in RELION 4.0, and after import into M, particle pose refinement, image warp grid (8×8 resolution), and stage angles were refined for three iterations. For all M refinement steps, 3 sub-iterations were carried out and 40% of available resolution was used for the first sub-iteration.

The subtomogram averaging pipeline for LSTc-bound HA samples was largely similar to HA alone. However, different masks were used at each stage to include density for neighbouring HA trimers instead of just the central HA trimer. Resolution of all final maps was estimated using the criterion of FSC=0.143 for masked and unmasked maps (40, 41). Local resolution range of maps were estimated using M (18).

To generate models, crystal structures of PR8 HA (PDB: 1RU7) and PR8 HA bound to LSTc (PDB: 1RVZ) was rigid body docked into our HA and HA-LSTc maps respectively using UCSF ChimeraX (30). Post-rigid body fitting, real space refinement was conducted using the Phenix software package (29).

### Visualization

3dmod was used to generate snapshots and movies of tomograms (33). UCSF ChimeraX (30) was used to visualize tomograms, subtomogram averages. PyMOL was used to visualize HA-HA interfaces (42). Seaborn and Matplotlib libraries were used to generate histograms (43, 44).

### Code and data availability

Custom Python scripts used for metadata conversion, starfile generation, inter-glycoprotein spacing calculation, and virion surface model generation are publicly available on Github (https://github.com/jqyhuang/influenza-analysis). Subtomogram averages and representative tomograms are deposited at the EMDB under the accession numbers EMD-XXXXX, EMD-XXXXX, EMD-XXXXX, EMD-XXXXX, EMD-XXXXX, EMD-XXXXX, EMD-XXXXX. Raw tiltseries are deposited at EMPIAR under the accession number EMPIAR-XXXXX.

## ACKNOWLEDGMENTS

The authors would like to acknowledge helpful discussions with the members of the Kelch, Munro, Grigorieff, and Schiffer labs, especially Matthew Unger, Clint McDaniel, and Drs Sabihur Rahman Farooqui and Nese Kurt Yilmaz. We would also like to thank the UMass Chan cryoEM Core facility for their help with data acquisition and for providing us with support and advice. We would like to acknowledge Life Science Editors for editing the manuscript. This work was supported by R01 GM143773.

## AUTHOR CONTRIBUTIONS

Conceptualization, Q.J.H, J.B.M, C.A.S, and M.S.; Methodology, Q.J.H, N.G, J.B.M, C.A.S, and M.S.; Software, Q.J.H; Formal Analysis, Q.J.H, R.K.; Investigation, Q.J.H, R.K, K.S.; Visualization, Q.J.H, Writing – Original Draft, Q.J.H Writing – Review & Editing, Q.J.H, R.K, K.S, N.G, J.B.M, C.A.S, and M.S;Funding Acquisition, J.M.B. and M.S.; Supervision, C.A.S, and M.S. C.A.S and M.S. contributed equally in project conceptualization, manuscript writing, and supervision. All authors approved of the final manuscript.

