## Supplemental Materials for "Virion-associated influenza hemagglutinin clusters upon sialic acid binding visualized by cryo-electron tomography"

Fig. S1. CryoET neural network analysis workflow. This workflow depicts the data analysis process to quantify virion morphology measurements and for particle picking of PR8 HA.

Fig. S2. General subtomogram averaging workflow for initial orientation estimation for HA and HA-LSTc subtomograms.

Fig. S3. Subtomogram averaging workflow for HA.

Fig. S4. Subtomogram averaging workflow for HA-LSTc.

Fig. S5. Inter-glycoprotein distances for HA and HA-LSTc. Box plots of inter-glycoprotein distance measured between central glycoprotein illustrated in Fig 2 and its closest two HA neighbours. Top and bottom lines indicate the interquartile range and the middle line median. Whiskers plot 5-95% of values.

Fig. S6. 2D classification of trimeric HA-LSTc array.

Fig. S7. Resolution estimates for subtomogram averages. Local resolution estimation and Fourier Shell Correlations (FSCs) of unmasked and masked maps for HA (a-b); HA-LSTc (100 μM) pair I (c-d); HA-LSTc (6.5 mM) central trimer (e-f); HA-LSTc (6.5 mM) pair I (g-h), and HA-LSTc (6.5 mM) pair II (i-j). Reported resolutions were measured at FSC=0.143

Table S1. Summary table of tilt-series collection

| **Sample** | **PR8** | **PR8 + 6.5 mM LSTc** | **PR8 + 100 μM LSTc** |
| --- | --- | --- | --- |
| **Microscope** | Titan Krios | | |
| **Voltage** | 300 kV | | |
| **Detector** | K3 | | |
| **Tilt range** | 0 to ±66° | 0 to ±51° | 0 to ±66° |
| **Tilt step** | 2 | 3 | 2 |
| **# subframes** | 4 | 6 | 4 |
| **Defocus range** | 4-8 µm | | |
| **Total Dose** | 120 | | |
| **Pixel size (A)** | 2.09 | | |

Table S2. Summary table of subtomogram averages.

|  | **Unliganded HA** | **Central HA from trio of HAs (PR8 + 6.5 mM LSTc)** | **Pair I from trio of HAs (PR8 + 6.5 mM LSTc)** | **Pair II from trio of HAs (PR8 + 6.5 mM LSTc)** | **Pair I from trio of HAs (PR8 + 100 μM LSTc)** |
| --- | --- | --- | --- | --- | --- |
| **# of final subtomograms** | 12,063 | 17,266 | 12,379 | 10,548 | 9,283 |
| **Map resolution (FSC=0.143)** | 8.9 Å | 8.5 Å | 8.9 Å | 10 Å | 9 Å |
| **EMDB ID** | EMD-XXXXX | EMD-XXXXX | EMD-XXXXX | EMD-XXXXX | EMD-XXXXX |

Movie S1. Representative tomogram of PR8. Tomogram corresponds to the slice shown in Figure 1A. Scale bar = 50 nm.

Movie S2. Representative tomogram of PR8. Tomogram corresponds to the slice shown in Figure 1B. Scale bar = 50 nm.

Movie S3. Initial HA model in comparison with real space refined model. Morph movie between rigid body docked crystal structure and flexibly fitted coordinates.

**Movie S4. Initial HA-LSTc model in comparison with real space refined model.** Morph movie between rigid body docked crystal structure and flexibly fitted coordinates.
